# 3-Dimensional Magnetic Resonance Imaging of the Freely Moving Human Eye

**DOI:** 10.1101/2020.06.26.172791

**Authors:** Benedetta Franceschiello, Lorenzo Di Sopra, Astrid Minier, Silvio Ionta, David Zeugin, Michael P. Notter, Jessica A.M. Bastiaansen, João Jorge, Jérôme Yerly, Matthias Stuber, Micah M. Murray

**Affiliations:** The Laboratory for Investigative Neurophysiology (The LINE), Department of Radiology, University Hospital Center and University of Lausanne, Lausanne, Switzerland; Department of Ophthalmology, University of Lausanne, Jules Gonin Eye Hospital, Fondation Asile des aveugles, Lausanne, Switzerland; Department of Radiology, University Hospital Center and University of Lausanne, Lausanne, Switzerland; École Polytechnique Fédérale de Lausanne (EPFL) Lausanne, Switzerland; Center for Biomedical Imaging (CIBM), Lausanne, Switzerland; Department of Hearing and Speech Sciences, Vanderbilt University Nashville, TN, USA

**Keywords:** magnetic resonance imaging, vision, compressed sensing, eye

## Abstract

**Abstract:** Eye motion is a major confound for magnetic resonance imaging (MRI) in neuroscience or ophthalmology. Currently, solutions toward eye stabilisation include participants fixating or administration of paralytics/anaesthetics. We developed a novel MRI protocol for acquiring 3-dimensional images while the eye freely moves. Eye motion serves as the basis for image reconstruction, rather than an impediment. We fully reconstruct videos of the moving eye and head. We quantitatively validate data quality with millimetre resolution in two ways for individual participants. First, eye position based on reconstructed images correlated with simultaneous eye-tracking. Second, the reconstructed images preserve anatomical properties; the eye’s axial length measured from MRI images matched that obtained with ocular biometry. The technique operates on a standard clinical setup, without necessitating specialized hardware, facilitating wide deployment. In clinical practice, we anticipate that this may help reduce burden on both patients and infrastructure, by integrating multiple varieties of assessments into a single comprehensive session. More generally, our protocol is a harbinger for removing the necessity of fixation, thereby opening new opportunities for ethologically-valid, naturalistic paradigms, the inclusion of populations typically unable to stably fixate, and increased translational research such as in awake animals whose eye movements constitute an accessible behavioural readout.

**Author contributions:** B.F., L.D.S., M.S., and M.M.M. conceptualised the problem. B.F. and L.D.S. developed, implemented, and tested the protocol. A.M. provided optometry assessments and assisted with eye movement analysis. S.I., D.Z., and M.P.N. assisted with installation of the eye-tracking system within the MRI scanner. J.A.M.B., J.J. and J.Y. contributed with the MRI sequences and compressed sensing framework. B.F., L.D.S. and M.M.M. drafted the manuscript, and all authors contributed to internal review.

**Competing interests:** B.F., L.D.S., J.A.M.B., J.Y., M.S., and M.M.M. declare the following competing financial interest: a patent application for the protocol described in this manuscript has been filed (patent application: EP19160832). A.M., S.I., D.Z., M.P.N. and J.J. declare no competing financial interests.

## Introduction

Despite the near-ubiquity of magnetic resonance imaging (MRI) techniques in clinical practice and research, its application in the field of ophthalmology poses major challenges due to the presence of eye movements and the reduced field of view of the image. Eye movements introduce motion artefacts and preclude the applicability of standardized procedures in clinical practice. The reduced field of view translates into a low signal-to-noise ratio and hence requires fine optimization of the contrast and pulse sequences. Other imaging methods have been introduced as screening technologies in ophthalmology, principally using visible or near-infrared light. Among those employed in anatomical imaging, confocal imaging (Webb, 1996), optical coherence tomography (OCT) (Fujimoto et al., 2016) and Fundus Photography (Panwar et al., 2016) are among the most widespread. They provide high-resolution images of the retina and the eye, both from the structural and angiographic perspectives. Such notwithstanding, disadvantages of these methods include stationarity of the eye during data acquisition, dependence on the clarity of the outer eye (i.e. lens, cornea, and vitreous humour), and most importantly an unobstructed pathway of light from the cornea through the lens and the retina. These techniques are otherwise inapplicable, such as when calcification is present or when there are changes in the normal tissues (e.g. in the case of tumours).

MRI provides a characterization of the tissues based on the nuclear magnetic resonance signals emitted by targeted nuclei, readily allowing for the differentiation between tissues and structures as well as between healthy and pathological states. Eye structures, such as the retina, iris, ciliary body, lens and aqueous and vitreous humours as well as surrounding tissues, such as muscles, in humans or animals can be visualized in vivo (Fanea and Fagan, 2012), reaching the retina resolution in animals (Cheng et al., 2006; Shen et al., 2006). Anatomic information coming from T2-weigthed or T1-weighted mapping can identify retinal detachment (Deans et al., 1988) and retinal abnormalities (Bahn et al., 1998), as well as variations in eye volume, surface area and shape in animals (Tkatchenko et al., 2010) and humans (Lim et al., 2011).

Proofs of concept of the feasibility of functional magnetic resonance imaging (fMRI) of the retina have been conducted in animal models based on blood flow measurement (Li et al., 2008; Muir and Duong, 2011), blood volume measurement (Nair et al., 2011; Shih et al., 2011), and blood oxygen-level dependent (BOLD) MRI during physiological challenges (Cheng et al., 2006) and visual stimuli (De La Garza et al., 2011; Duong et al., 2002).

However, the extension of extant protocols to humans remains challenging. High-resolution anatomic imaging of the human eye has been reported (Richdale et al., 2009), while a first attempt to BOLD MRI during a gas challenge has been reported (Zhang et al., 2011b). However, the proposed fixation protocol for blinking control introduces an important confound, as it affects the overall statistics estimation of the BOLD activation, and blinking itself has been shown to produce BOLD signal changes in the brain (Hupé et al., 2012). In light of such, there is a growing recognition of how MRI techniques can be applied in ophthalmology, including in clinical practice (Fanea and Fagan, 2012; Townsend et al., 2008). Nowadays the use of MRI of the eye in clinical practice is limited to the anatomical imaging of sedated or anesthetized subjects for retinoblastoma diagnosis, because tumour tissue often has unique features that are identifiable through MRI and because it is important to ascertain if tumours have metastasized into the brain (Moulin et al., 2015; Sirin et al., 2016; Townsend et al., 2008).

MRI has the potential to overcome the limitations coming from other screening technologies in ophthalmology, and the current findings in animal models also suggest that MRI could introduce new, and potentially more comprehensive, measures of the eye’s structure as well as function, extending across the entire eye-brain circuitry. However, dedicated sequences and techniques need to be developed so that MRI scanners can be fully exploited for eye-brain imaging, taking advantage of its extensive presence in healthcare facilities. Eye movements remain a major problem that technological advances in the field confront. MRI acquisition time is slow compared to the speed of drifts, tremors, saccades, and blinks of the eye; movements that are particularly pronounced when subjects are asked to stay still in the scanner and maintain fixation (Duong, 2014; Fanea and Fagan, 2012; Townsend et al., 2008). To our knowledge the first and only attempt to solve motion in the eye has been proposed by Sengupta et al. (Sengupta et al., 2017). However and in contrast to the methods that will be introduced here, their technique only allowed for the acquisition of 2-dimensional images, eye movements were not controlled nor monitored externally, and time-resolved image reconstruction was performed instead of a position-based one, which minimises noise.

Inspired by the successful novelties introduced in motion-resolved 3D MR imaging of the heart (Coppo et al., 2015; Di Sopra et al., 2019; Feng et al., 2018), we ideated and validated an MRI acquisition framework that demonstrates the feasibility of motion-tracking and retrospective motion-resolved 3D isotropic image reconstruction of the eye’s dynamics, while also including the whole brain within the field of view. This technique overcomes the limitations typically caused by eye movements, first by inducing the eye to move in a guided way and second by using eye movements as extra-dimensions in a “compressed sensing” image reconstruction framework, instead of considering motion as an issue to correct during a post-processing step (**Figure 1**). To our knowledge, this is the first attempt in this direction capable of providing an isotropic 3D image of the eye and brain, while it freely moves under visual stimulation.

**Figure 1.**
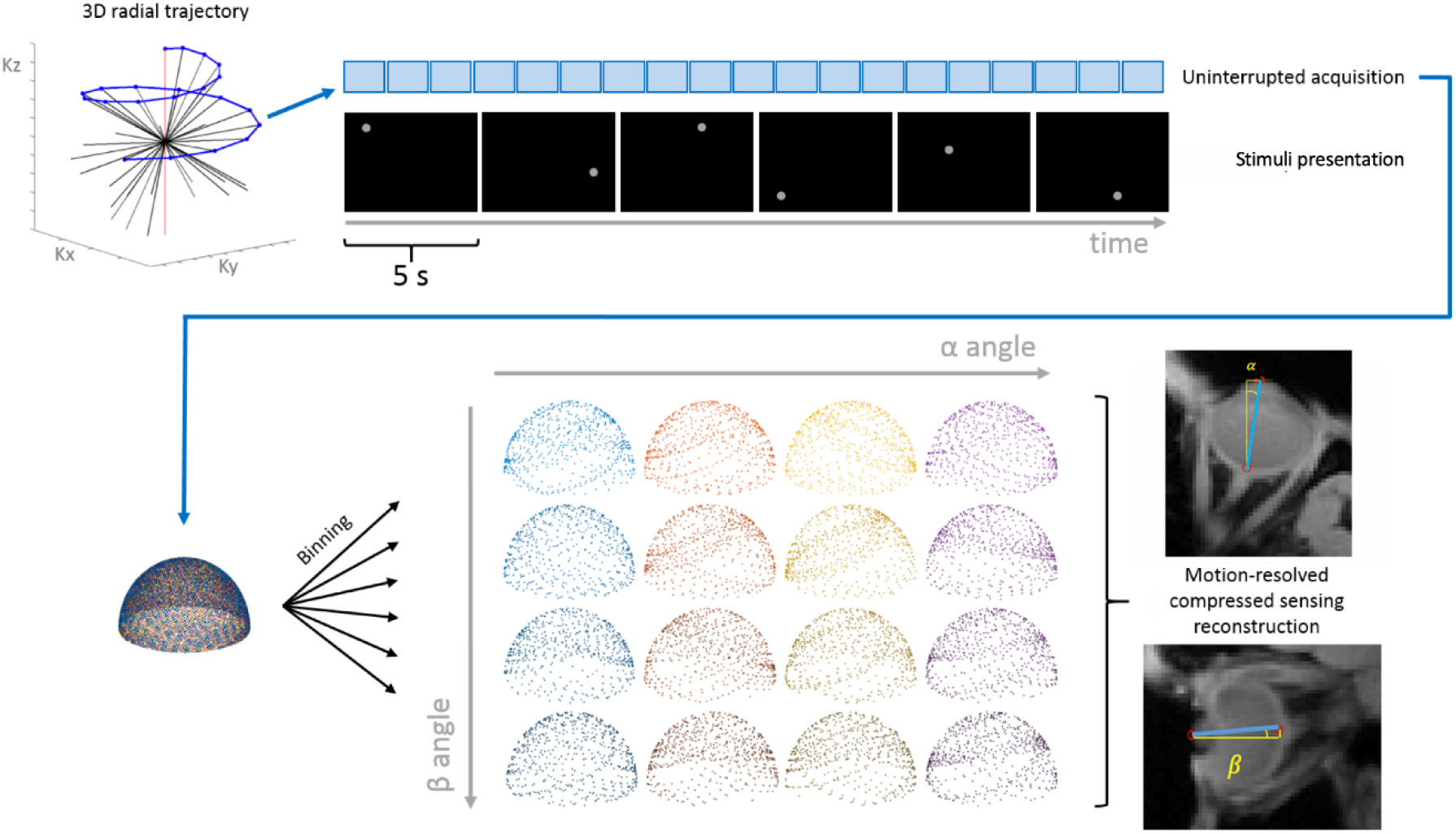
A schematic illustration of the framework from image acquisition to reconstruction. Each one of the 16 final motion-resolved images (associated to the locus of stimulus presentation) is obtained from an under-sampled set of about 5000 k-space readouts (i.e. 81906/16). Nevertheless, uniform k-space coverage regardless of the stimulus presentation order is ensured by the radial phyllotaxis trajectory properties. The stimulus presentation scheme was parametrized by the angles (α, β). The idea at the basis of the k-t sparse SENSE algorithm is that close-by positions (such as the blue and the red cloud of dots) share a high amount of information about the k-space, allowing to perform image reconstruction from an under-sampled acquisition of k-space lines.

## Material and Methods

Data were acquired from nine healthy adult volunteers (aged 27-34 years; 4 men and 5 women) on a 3T clinical MRI scanner (MAGNETOM Prisma^fit^, Siemens Healthcare AG) with a 22-channel head coil, using a prototype uninterrupted gradient recalled echo (GRE) sequence with lipid-insensitive binomial off-resonant RF excitation (LIBRE) for fat suppression (Bastiaansen and Stuber, 2018). The study protocol was approval by the cantonal ethics committee (protocol number 2018-00240). The acquisition used a 3D radial sampling pattern, the spiral phyllotaxis trajectory (Piccini et al., 2011), where each interleaf is rotated by the golden-angle to allow uniform k-space coverage. The continuously acquired data, as enabled by the free-running approach to data collection (Coppo et al., 2015), can be arbitrarily partitioned into different bins thanks to the golden-angle distribution properties. Eye-tracked trajectories, together with related trial number and temporal synchronization information, were extracted from the eye-tracking software (detailed below). The processed eye-tracker data were used to bin the time intervals of each motion state and to match the k-space readouts corresponding to the same stimulus presentation, hence leading to the same motion-resolved 3D image. In other words, eye-tracker trajectories were used to separate the k-space readouts into different volumes. Motion-resolved 5D image reconstruction (x-y-z-α-β dimensions, where α and β represent the eye angular rotations in the horizontal and vertical directions, respectively) was performed using a k-t sparse SENSE algorithm (image under-sampling 8.8%), exploiting sparsity both along the α and β directions. The values of α and β are deduced from the eye-tracker recordings and correspond to those determined from the reconstructed images by a trained optometrist (A.M.), once normalized. With this protocol, reconstruction required approximately 1 hour per subject, though this time is dependent on the available IT infrastructure.

The FoV was 192mm^3^ with 1mm^3^ isotropic resolution, TR/TE=6.4/2.94ms, receiver bandwidth BW=501 Hz/px, and radiofrequency excitation angle FA=5°. This encompassed the eyes as well as the majority of the head and brain. Eye movements were tracked using an eye-tracking system (EyeLink 1000Plus, SR Research) synchronized with the MRI scanner via Syncbox (NordicNeuroLab). An Experiment builder (EyeLink) program was developed and used to control the calibration of the Eye-Tracker from outside the scanner room and to correctly synchronise the different hardware components of the experiment. The right eye was the one tracked during the acquisition. Eye movement trajectories were recorded using infrared light, with a sampling rate of 2000Hz, through a mirror positioned inside the scanner bore, replacing the standard head-coil mirror usually available, which is not infrared compatible.

The stimulation protocol was divided into 3 distinct phases, all consisting of a grey circle positioned at specific locations on a black background. These circular stimuli guided the eye movements. First, an initial period of fixation was performed, where the image presented to the participant was the static grey circle positioned at the centre of the screen. This first part of the experiment allowed for performing the sequence localizer while the eye was in a static position. Second, 96 visual stimuli were presented to each participant. Each stimulus corresponded to one among 16 different locations the grey circle on a 4×4 grid. Each stimulus presentation had a duration of 5 seconds and was repeated 6 times at distinct and randomized moments during the experiment. This part of the acquisition lasted for 8 minutes in total. Third, the fixation circle was presented again, as in the first part of the experiment, to conclude the acquisition. The presentations during the second phase of the experiment were opportunely randomized to ensure a uniform sampling distribution of the readouts in k-space during the following retrospective motion-resolved reconstruction step. A total of 81,906 readout profiles, divided into 3,723 interleaves, were acquired.

## Results

### Motion-resolved image reconstruction

As can be seen in **Figure 2** (see also Supplementary Information for movies), 3D motion-resolved images of the eye with 1mm^3^ isotropic resolution were successfully acquired and reconstructed, devoid of motion artefacts, in all participants. Moreover and as is best visible in the movies, there is excellent coherence between the motion resolved in the reconstructed images and the stimuli presented. On each trial, a grey disc on a black background was presented at one of sixteen pseudo-randomised positions on a hypothetical 4×4 grid that the subject followed with eyeball rotation both in the vertical and horizontal directions.

**Figure 2.**
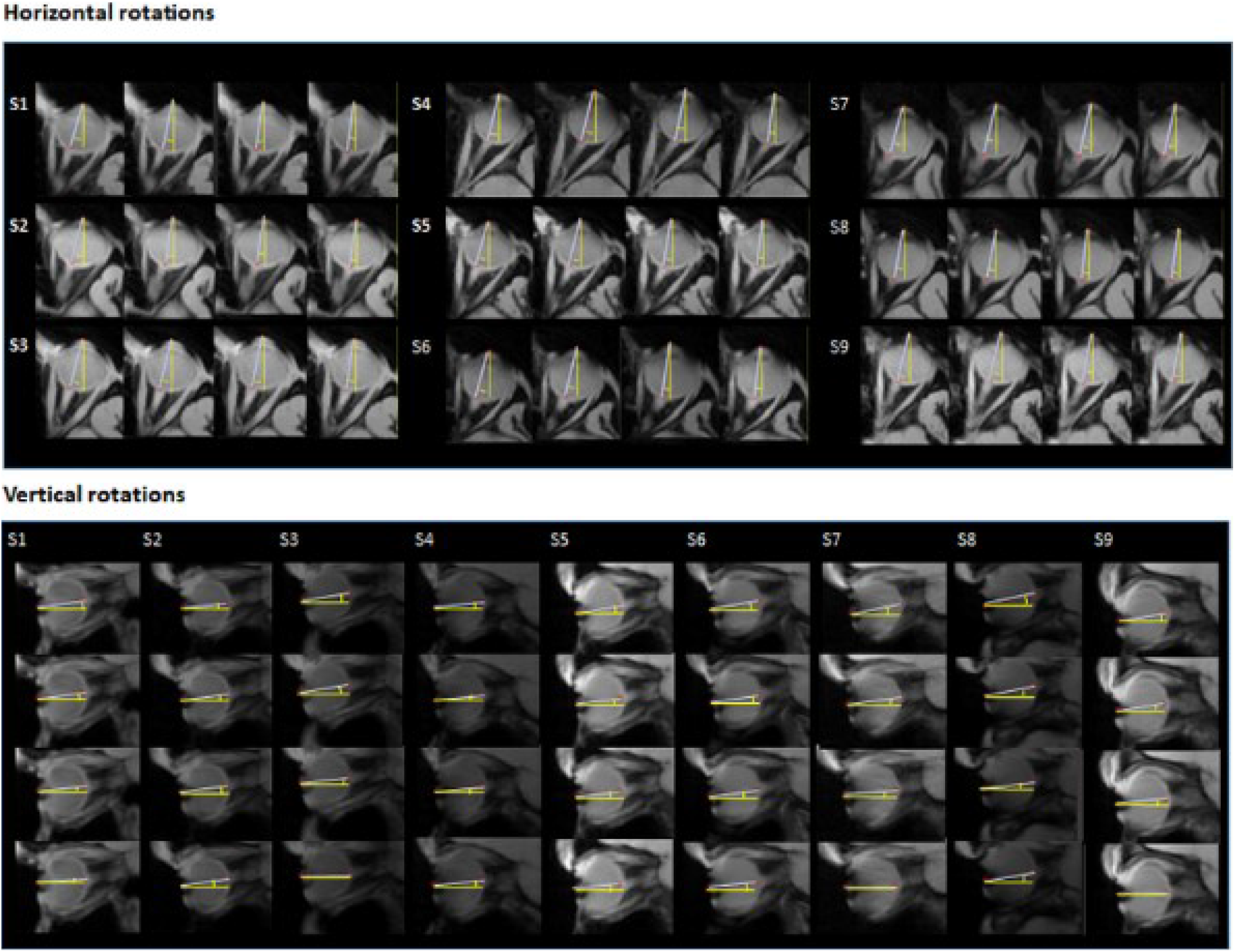
A series of reconstructed images based on horizontal and vertical rotation of the eye. For each subject (S1-S9) we were able to reconstruct artefact-free images of the eye (and brain), corresponding to a specific angular rotation of the eye in either the horizontal or vertical direction that follows each of the sixteen positions of the visual stimuli used here. Every position among the sixteen shown to every participant could be uniquely identified by the combination of horizontal and vertical angles of rotation of the eye, resulting from four total horizontal rotation angles and four total vertical rotation angles.

### Temporal image reconstruction

**Error! Reference source not found.** displays a comparison between our motion-resolved images (left side of figure) and those we would obtain when letting the eye freely move (right side of figure). In our framework, the latter is obtained by binning together temporally consecutive lines of the k-space, despite the different position of the eyes. Likewise, the compressed sensing reconstruction could not exploit sparsity along any dimension. This situation approximates motion and under-sampling artefacts one would obtain if the eyes would move within a standard non-compressed sensing 1mm^3^ isotropic T1-weighted GRE acquisition In particular, the noisy images on the right have been retrieved through a 4D reconstruction (no k-t sparse SENSE) having the time, *t*, as a fourth dimension, instead of performing a 5D k-t sparse SENSE reconstruction. The sections in **Error! Reference source not found.**, right, are composed by readouts acquired continuously for 30s, matching the bin size of those volumes reconstructed with the technique proposed in this work.

### Quantitative assessment of motion tracking

To evaluate the performance of the reconstructed images in terms of the resolution of eyeball rotation, benchmarks defining the eyeball axis were manually determined following guidelines from an experienced optometrist (A.M.). The vertical and horizontal rotation axes were manually determined, and the corresponding rotation angles were computed as:

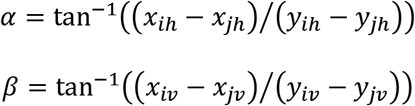

where *i* index denotes the optic nerve, *j* the pupil, *h* refers to the horizontal plane and *v* refers to the vertical plane. The pupil and optic nerve benchmarks were extracted from two different slices, as they were usually not belonging to the same sagittal slice (due to the horizontal eyeball rotation). Furthermore, the vertical and horizontal rotations induced by the experimental setup (theoretical angle) were considered to estimate the eyeball rotation angles as resolved by the reconstructed images in response to the stimuli presented (measured angle). Both measures were normalized, because they come from different reference frames. More specifically, the theoretical angle corresponds to a behavioral rotation of the eye, while the measured angle corresponds to the anatomical rotation. This latter measurement is expected to be proportional to the rotations induced by the stimulation and to the one recorded by the eye-tracker, the last ones used to reconstruct the volumes. To summarise, the eye-tracker trajectories were used to reconstruct the volumes corresponding to different positions of the eyeballs, while the target position of the visual stimuli were used to compare the rotation extracted by the optometrist from the MR-images.

The projector screen was located 102cm from the head coil, and the projected grey dots were positioned on a 4×4 grid with an average horizontal distance of 8.13cm and a vertical one of 6.2cm. This translates to an expected average rotation of the eyeball of about 4.55° in the horizontal direction and 3.47° in the vertical direction of motion. Among the 16 presentation positions, the top-left one was selected as the reference angle; and the difference with the other 15 positions was calculated. This difference allows for determining how well the proposed technique enables the correct estimation, from the motion-resolved reconstructed images, of the eyeball angular rotation during stimulation. For each participant and each direction of rotation, we calculated non-parametric correlation between the estimated rotation in the reconstructed MRI volumes and the rotation of the eye measured by the eye-tracker using Spearman’s rho (**scatterplots in Figure 4**). In the horizontal direction, these values ranged from 0.92 to 0.97 across participants and were each statistically reliable (p<0.001 in each case). In the vertical direction, these values ranged from 0.86 to 0.98 across participants and were each statistically reliable (p<0.001 in each case). These results indicate the capacity of our approach to track successfully both horizontal and vertical eye motion at an imaging level.

**Figure 3.**
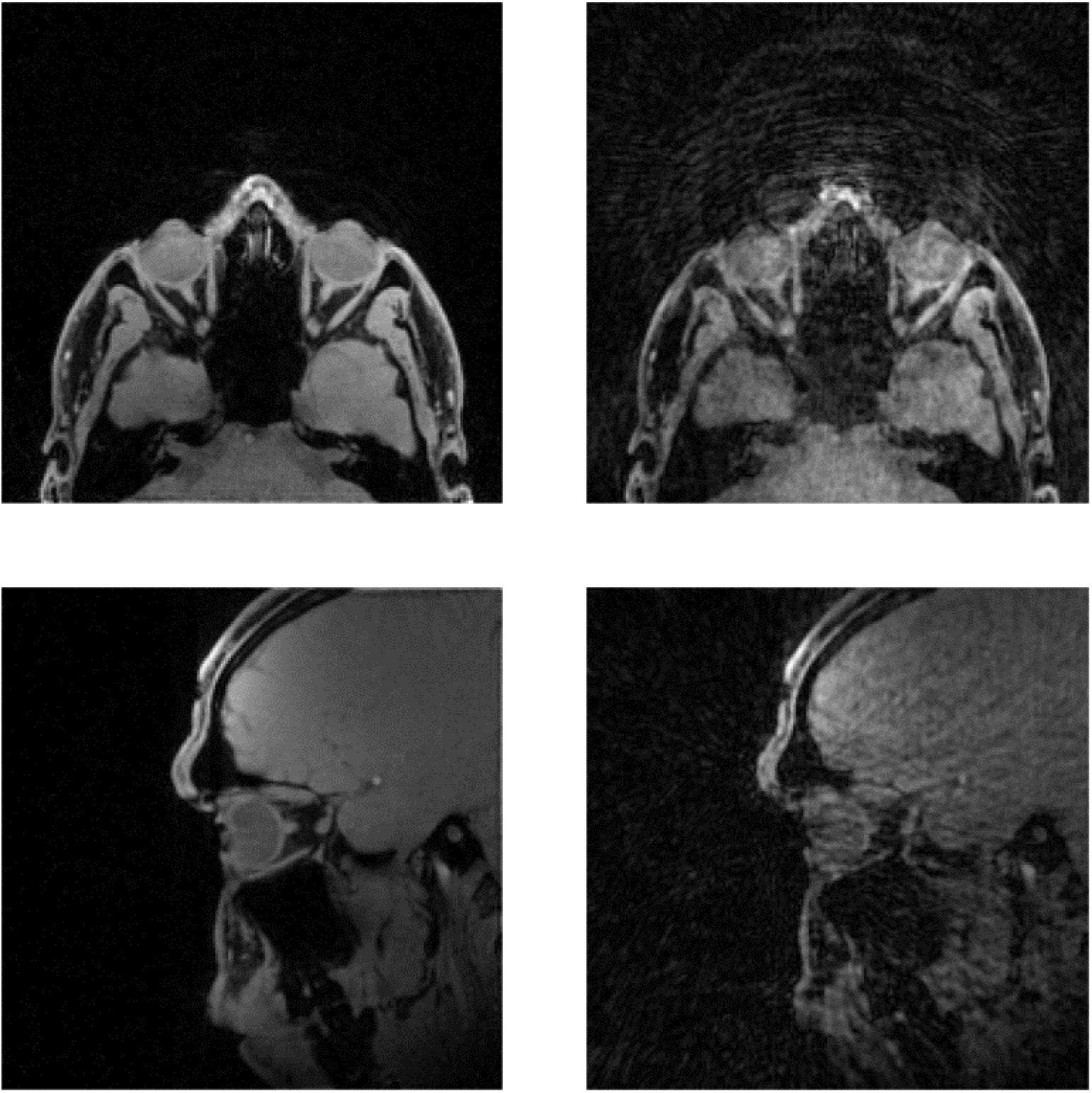
Comparison of images when binned based on eye position versus when binned consecutively. (Left). Standard volume reconstructed through our technique. On top the axial slice is presented, and on the bottom a sagittal view. (Right) When volumes are binned together based on continuously acquired k-space readouts motion artefacts are apparrent, as information corresponding to different positions assumed across time by the eyes is pooled together. This situation approximates the classical motion and under-sampling artefacts we would obtain while acquiring a T1-weighted image and reconstructing it without a motion-resolved compressed sensing algorithm.

**Figure 4.**
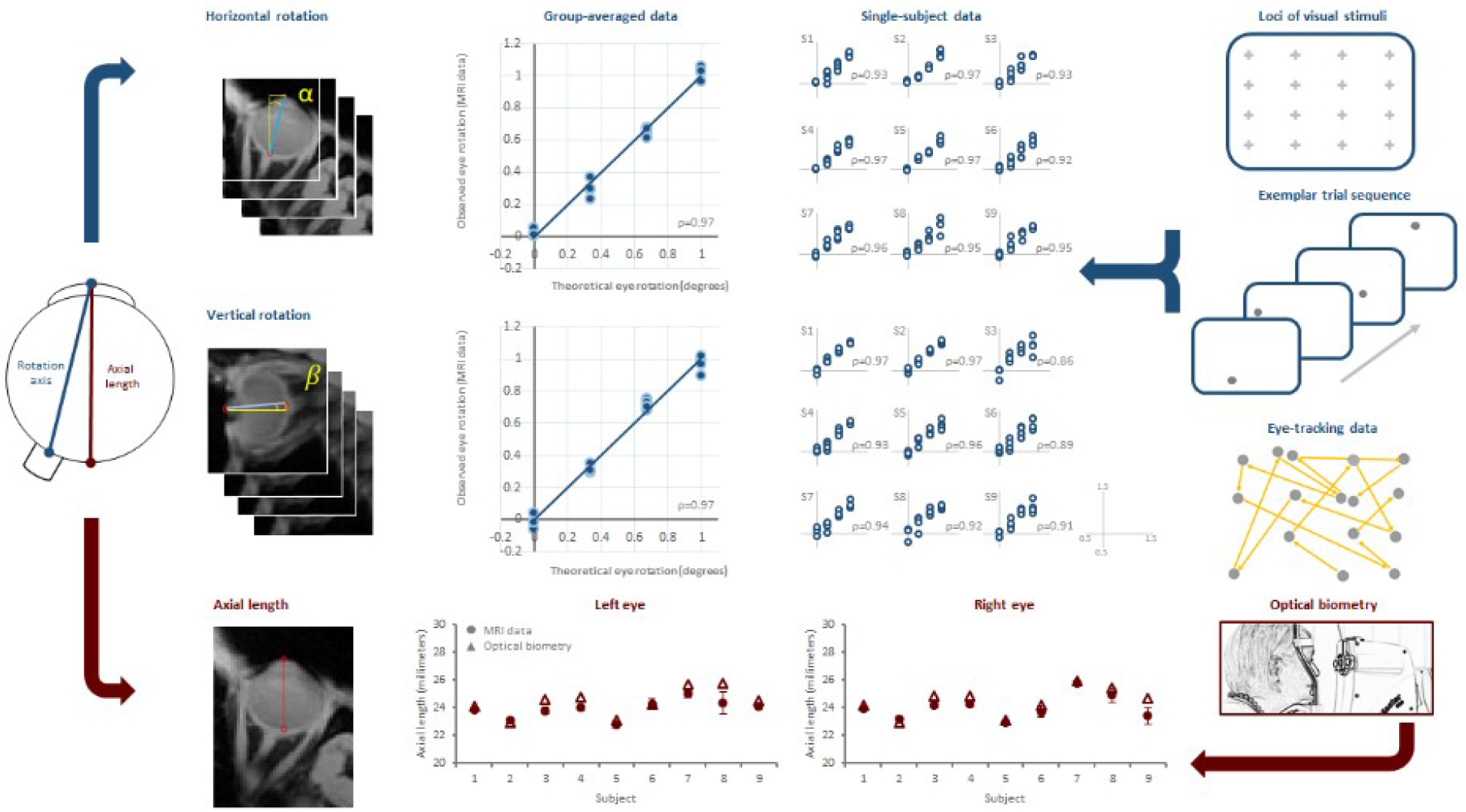
Quantitative validation of image quality. Two classical benchmarking techniques were used to quantitatively assess not only the resolution of the images obtained from the protocol, but also their informativeness. First, the eye-tracker assessed motion changes (blue). Second, optical biometry evaluated the axial length of the eye (red). In the top part of the image we observe how the α, β –rotation angles estimated from the MR-images, linearly correlate (at a mean and individual subject level; Spearman’s rho indicated) with the theoretical rotation performed by the eye during the visual stimulation (both axes represent normalised differences in degrees). In the bottom part of the figure, it is shown how the axial length estimated across all presentations from the MR-images has the same order of magnitude (mm) of those estimated by the biometry (μm accuracy).

### Quantitative assessment of eye image quality

Next, we assessed the extent to which our approach yields comparable metrics to those obtained in standard clinical practice. Optical biometry is a non-invasive clinical exam based on interferometry used to enhance anatomical structures of the eye. A partially coherent infrared light source is emitted toward the eye where it undergoes various reflections to form an emerging signal. The reflected interferences are used to calculate the optical distances that separate the different ocular structures (Santodomingo-Rubido et al., 2002). All of the MRI participants underwent an ocular biometry (IOLMaster 700, Zeiss) to enable us to estimate the axial length of each eye (Lam et al., 2001). In clinical practice, axial length is a principle diagnostic tool for myopia. We also estimated axial length from the reconstructed MRI data (performed by a trained optometrist). The lower portion of **Figure 4** displays these measurements for each participant and eye. The mean difference (±standard deviation) between these measures was 0.48±0.34mm for the right eye and 0.55±0.43mm for the left eye. Across individuals, the MRI-based and biometric estimates of the eye’s axial length were strongly correlated for each eye (Spearman’s rho = 0.92 and 0.77 for the right and left eye, respectively; p<0.016 in both cases). This pattern undoubtedly proves that the technique, once optimised, could provide information about anatomical structures consistent with those obtained through classical optometric methods.

## Discussion

We demonstrated the feasibility and accuracy, in terms of image quality and replication of ophthalmic measurements, of uninterrupted acquisition of the eye while it moves under visual stimulation. Instead of eye motion being a nuisance, we show how to use it as the basis for image reconstruction. This was made possible by a compressed sensing framework from which 5-dimensional images are retrospectively reconstructed based on eye position. Here, this motion-resolved imaging was based on horizontal or vertical position. However, in principle this can be extended to any formulation of the eye’s rotation and its sequence over time. In the present case, we used a relatively long fixation duration. However, this can readily be shortened to more naturalistic durations, as the only *a priori* limit based on the MR physics of our protocol is the acquisition time of a single k-space radial readout (currently TR=6.4ms). The method detailed here has tremendous opportunity for further improvement of the contrast and spatial resolution of the acquired images by finding and tuning the most suitable parameters for eye-brain MRI protocols with uninterrupted acquisition sequences and retrospective motion-resolved reconstruction. This optimization is required both for technical and research-related purposes. However, the versatility of the approach renders it easily adaptable and targeted. Not only were reconstructed images reliably following eye position, but we also proved that the obtained results are consistent with classical anatomical estimates of the eye (axial length measures) obtained through the state-of-the-art of clinical ophthalmic evaluations. More generally, the methods introduced here can be applied in situations both when the objective is to resolve eye motion itself and when the objective is to avoid confounds produced by eye motion. That is, the user can reconstruct high-resolution images of the eye and brain either throughout the trajectory of the eye or for specific positions (e.g. exclusively when fixation was central, at a given eccentric location, etc.). Consequently, we believe that the current methods and the enhancements that are readily implemented are harbingers of a fundamental paradigm shift in how MRI research is applied in ophthalmology and brain imaging alike.

The protocol introduced here has wider implications, particularly in the domain of brain imaging and cognitive neuroscience. Current MRI-based neuroscience techniques typically necessitate participants’ fixation (Gitelman et al., 2000; Warnking et al., 2002), thereby limiting the varieties of paradigms that can be used and their real-world comparability. Central fixation is typically considered as a control not only of what image is projected onto the retina and therefore into the brain, but also of a participant’s focus of attention. However, central fixation is arguably neither representative of naturalistic viewing (and by extension perception), nor is it forcibly indicative of participants’ attention. Some data even suggest that activations observed during central fixation tasks within the frontal eye fields and superior parietal lobule are more the product of suppressing eye movements than of attentional load (Culham et al., 2001). These aspects have contributed to an interest in studying more naturalistic paradigms and in understanding perception as a consequence of active sensing (Schroeder et al., 2010). Although data acquisition is possible during free viewing, provided that eye-position is tracked (Kanowski et al., 2007; Kimmig et al., 1999), image reconstruction, and thus data analysis, typically excludes the eyes and is limited to the brain. The present results highlight a roadmap for fast and fixation-unconstrained MR imaging that would permit investigations along the full eye-brain circuitry. Our technique allows for image reconstruction as a function of eye position and thus would not be limited to only when central fixation is achieved. Instead and as the present study shows, MR images are binned according to eye position. This would allow for more naturalistic paradigms as well as the ability to examine the interplay between scanning/fixation behaviour, eye-brain activity and perception. For example, hippocampal activity in humans correlates with the number of fixations during the encoding of novel faces (Liu et al., 2017); a relationship that declines with aging (Liu et al., 2018). Eye position is known to not only affect visual processing, but also auditory responses throughout subcortical and cortical stages of hearing (Groh et al., 2001; Werner-reiss et al., 2003). Moreover, there is ample evidence for both head-centered and eye-centered spatial representations across multiple sensory modalities (Mullette-Gillman et al., 2005). Our methods provide a means to accessing these topics via MRI, while balancing naturalistic behaviour of participants with fine-grained experimenter control over analyses. More generally, the approach we describe in our study would, in turn, facilitate MR-based investigations of notions regarding active sensing (Schroeder et al., 2010; van Atteveldt et al., 2014) or expertise (Quiroga and Pedreira, 2011) and their inter-individual variability.

Moreover, the necessity to fixate also restricts the range of compliant participants; something often challenging for paediatric and aged populations alike (Greene et al., 2018; Plank et al., 2017). As such and because we used standard MRI equipment and eye-tracking hardware, the approach we have developed and validated is readily deployed to the broad scientific community and impacts not only the breadth of participant inclusion, but also the extent of naturalistic paradigms that can be investigated (Nishimoto et al., 2017). In particular, infants/children and the elderly are two such populations where fixation is particularly difficult to maintain (e.g (Byars et al., 2002) for a review of the challenges of fMRI in 5-18 year-olds). In a future in which the orientation of the eyeball could be extracted from the images in an automatic way during acquisition, one could for example envision a closed-loop system wherein the image position adapts to eye position such that it is always projected onto the fovea. This would remove the onus of maintaining central fixation (see e.g (Son et al., 2020) for a very recent article in this direction). Furthermore, this would apply both to research in humans, but also to research in animal models. In the latter case, there is added translational value when eye fixation and anaesthetics can be minimised (if not rendered unnecessary) and when the experimenter can use eye movements as a behavioural readout (Meyer et al., 2018). The renewed interest in characterizing perceptual processes within an active sensing framework underscores the need to allow for unrestrained eye movement in experimental protocols (Barczak et al., 2019; Schroeder et al., 2010). A further advantage of circumventing the need for an eye-tracker is that eye position can be followed even when the eyes are closed, when eye position cannot be readily controlled, or when the pupil is obscured (and therefore unsuitable for typical eye-tracking devices). This would open new possibilities in fields such as sleep and dream research (Desseilles et al., 2011; Siclari et al., 2017), mental imagery (Fourtassi et al., 2017), as well as more generally for understanding brain activity in disorders of consciousness (Owen, 2013).

Importantly, our method could in principle extend to the physiological interactions between the eyes and other signals, such as heartbeat, respiration, etc. It therefore constitutes a breakthrough for what concerns methodology in neuroscience, as physiological signals (ocular, respiratory, cardiac, etc.) could theoretically serve as the bases for image reconstruction of images, avoiding correlation analysis between images and signals at a post-processing level (Chen et al., 2019). This integration would provide a potential account for how our eye muscles and brain functionality interact with the main body regulatory systems; a domain that has thus far remained underexplored. In these ways, our protocol provides a clear roadmap for comprehensive eye-to-brain measures in terms of both structure and function.

From the clinical standpoint, the benefits of being able to accurately image a freely moving eye are clear. Patients need no longer fixate or even be capable of such (Henderson and Choi, 2015; Kimmig et al., 1999). An important next step will be the evaluation of the efficacy of the developed technique in targeted clinical populations. In terms of feasibility, the duration of the current protocol is relatively short (approximately 8 minutes) and can undoubtedly be further optimised. Duration notwithstanding, the possibility of reducing or eliminating the need for administering paralytics or anaesthetics during MRI acquisition is highly desirable both for patient health and in terms of personnel and infrastructural costs for the hospital (Duong, 2014; Duong and Muir, 2009; Moulin et al., 2015). MRI provides anatomical and functional information that cannot be explored with other classical ophthalmologic techniques (OCTs, biometry, etc.), such as those concerning eye-dynamics, optic nerve anatomy, whole eye-brain pathway. Hitherto, motion has constituted the main reason why there are no studies presenting functional activation of the neural retina after visual stimulation in humans (Duong, 2014; Duong et al., 2002)(Zhang et al., 2011a, 2011b). We believe our approach could open a whole range of applications in clinical as well as fundamental visual neuroscience research, allowing for the introduction of new benchmarks and measures such as the anatomical assessment of eye movements, which might constitute an important biomarker for cognitive disorders (Anderson and MacAskill, 2013; Colligris et al., 2018; Fukushima et al., 2017; Liao et al., 2018). The methods introduced here demonstrate what is currently feasible and are certainly a harbinger of continued transformative developments.

## Acknowledgments

Financial support for this work has been provided by the Fondation Asile des aveugles (grant #232933 to M.M.M.), a grantor advised by Carigest SA (#232920 to M.M.M.), as well as the Swiss National Science Foundation (grants #169206 to M.M.M., #173129, #150828, and #143923 to M.S., PZ00P3_167871 to J.A.M.B., and PP00P1_170506 to S.I.).

**Supplementary Movies 1 and 2** These animated gifs show the series of image reconstructions as a function of eye movement in both the horizontal and vertical directions as the participant fixates on the locus of the visual stimulus. The viewer should note the field of view covers not only the eye but also the brain, and image reconstruction is achieved across the entire field of view.

